# Evaluating the therapeutic efficacy of the US FDA-designated ‘Orphan drug’, uttroside B in impeding hepatocarcinogenesis via induction of immunogenic apoptosis, using a murine model of Aflatoxin B1-induced liver carcinogenesis

**DOI:** 10.1101/2025.09.08.674785

**Authors:** Mundanattu Swetha, Chenicheri Kizhakkeveettil Keerthana, Tennyson P Rayginia, Kalimuthu Kalishwaralal, Sadiq C Shifana, Sreekumar U Aiswarya, Lekshmi R Nath, Sanjay Suresh Varma, V Arun, S Jannet, J Shirly, JS Aparna, Sankar Sundaram, Nikhil P Anto, Noah Isakov, Ravi S. Lankalapalli, Kuzhuvelil B Harikumar, Ruby John Anto

**Affiliations:** Division of Cancer Research, BRIC-Rajiv Gandhi Centre for Biotechnology, Thiruvananthapuram 695014, Kerala, India; Molecular Bioassay Laboratory, Institute of Advanced Virology, Bio 360 Life Sciences Park, Thonnakkal, Thiruvananthapuram, Kerala, India; Chemical Sciences and Technology Division, CSIR-National Institute for Interdisciplinary Science and Technology, Thiruvananthapuram 695019, Kerala, India; Department of Pathology, Government Medical College, Kottayam 686008, Kerala, India; The Shraga Segal Department of Microbiology, Immunology and Genetics, Faculty of Health Sciences, Ben-Gurion University of the Negev, P.O. Box 653, Beer Sheva 84105, Israel; Centre of Excellence in Nutraceuticals, KSCSTE, Government of Kerala, Thiruvananthapuram, 695317, Kerala, India

**Keywords:** Uttroside B, Aflatoxin B1, hepatocellular carcinoma, DNA adduct, immunogenic apoptosis, mutagenesis

## Abstract

The sustained exposure to aflatoxin B1 (AFB1), a mycotoxin produced by *Aspergillus sp*., is one of the fundamental causes of hepatocellular carcinoma (HCC). We have previously documented the exceptional anti-HCC potential and pharmacological safety of uttroside B, a phytosaponin isolated in our lab (Utt-B). The current results indicate that Utt-B mitigates tumor development in mice that have been subjected to AFB1 exposure. Utt-B was found to be cytotoxic towards primary liver cancer cells cultured from mice bearing AFB1-induced liver tumors and the compound effectively prevented the formation of AfBO-DNA adducts in AFB1-induced liver tumors as well as primary liver cancer cells. *In vitro* studies revealed that treatment with Utt-B resulted in the induction of damage associated molecular patterns such as, ROS, HSP70 and inflammatory cytokines, IL-1β and CXCL-10, suggesting the potential of Utt-B in triggering immunogenic cell death. Mechanistically, treatment with Utt-B enhances the antigen presentation potential, causes blockade of the major immune checkpoint molecules, namely, CTLA-4, PD-1, TIM-3, LAG-3 and TOX, and potentiates immunogenic apoptosis in the hepatic tumor microenvironment of mice pre-exposed to AFB1 via the activation of Zap70/Lck/GRZB signaling axis. Interestingly, it was also observed that Utt-B could abrogate the mutations induced by AFB1 in a concentration dependent manner. Taken together, the findings of the current study attest Utt-B as a propitious drug candidate against HCC.

## Introduction

Aflatoxin B1 (AFB1), a mycotoxin produced by *Aspergillus sp*., is a group 1 human carcinogen [1]. AFB1 contamination in food presents a substantial global health concern, and is one of the major causes of liver cancer, particularly in developing nations [2–4]. Aflatoxin contributes to about 4.6-28.2% of all annual hepatocellular carcinoma (HCC) cases worldwide [5]. Previous literature suggests that exposure to AFB1 induces severe immune suppression in the host [6, 7]. In the recent times, reprogramming of the tumor microenvironment (TME) to activate anti-tumor immune responses via induction of immunogenic cell death (ICD), a new type of regulated cell death has gained prominence [8]. Several chemotherapeutic agents such as, doxorubicin, oxaliplatin and paclitaxel induce ICD in tumor cells [9–11]. The chemotherapeutic effects of anti-cancer agents, coupled with their potential to induce ICD, culminates in sustained long-term therapeutic effects [12]. Certain classes of molecules known as the damage-associated molecular patterns (DAMPs), which are released from the dying tumor cells serve as molecular signals that induce the maturation and activation of dendritic cells (DCs) and the activation of cytotoxic T-cells, eventually leading to the induction of a robust anti-tumor immune response in the TME [13]. Previous reports from the lab provide compelling evidence on the exceptional chemotherapeutic efficacy of Utt-B against HCC [14–17].

The present study aims to evaluate whether the immunomodulatory effects of Utt-B complement the chemotherapeutic efficacy of the compound against Aflatoxin B1 (AFB1)-induced liver carcinogenesis.

## Materials and Methods

### Isolation and Purification of Utt-B

Isolation and purification of Utt-B from methanolic extract of *Solanum nigrum* Linn. leaves was done as previously reported [14].

### Cell lines

The liver cancer cell line HepG2 was purchased from ATCC. Mycoplasma tests were performed on parent cell lines and stable cell lines every 6 months.

### Chemicals and Antibodies

Cell culture reagents such as Dulbecco’s Modified Eagle Medium (DMEM) (GIBCO, 12800-017), Fetal Bovine Serum (10270-106), streptomycin sulfate (GIBCO, 11860-038), and Hoechst 33342 (H1399) were obtained from Invitrogen Corporation (Grand Island, USA). Poly Excel HRP/DAB detection system universal kit (PathnSitu Biotechnologies Pvt. Ltd, India, OSH001) was used for immunohistochemistry experiments. MTT reagent was purchased from TCI Chemicals (India) Pvt. Ltd (D0801). Antibodies against β-actin (12620S), Caspase 9 (9508S), PARP (9532S) and HSP70 (4876T) were obtained from Cell Signaling Technologies (Beverly, MA, USA) and PCNA (sc25280), Ki67 (sc23900), Pancytokeratin (sc8018) and AFP (sc130302) were purchased from Santa Cruz Biotechnology (Santa Cruz, CA, USA). Aflatoxin DNA adduct competitive ELISA kit (AKR-351) was purchased from Cell Biolabs Inc. DNA isolation kit for cells and tissues was procured from Roche (Cat. No. 11814770001). NucleoSpin RNA kit for RNA isolation was purchased from Macherey Nagel (740955.50). PowerSYBR Green PCR master mix (4367659) and High-capacity cDNA kit (4368814) were purchased from Applied Biosystems. Aflatoxin B1, tricaprylin and all other chemicals were purchased from Sigma Chemicals (St. Louis, MO, USA) unless otherwise mentioned.

### *In vivo* liver carcinogenesis model

The chemotherapeutic studies using Utt-B were carried out in DBA/2J mice, according to the pre-scribed guidelines, which were approved by the Institutional Animal Ethical Committee, RGCB (IAEC/536/RUBY/2016, IAEC/879/RUBY/2022). The DBA/2J strain of mice are susceptible to AFB1 and 90% of the mice develop HCC post-exposure to AFB1[18, 19]. Hepatic tumors were induced by oral administration of 6 µg/g body weight of AFB1 in tricaprylin in 3-week-old DBA/2J mice. The DBA/2J mice were randomly assigned to the three treatment groups: vehicle control (also mentioned as negative control in ‘results’, AFB1 alone (positive control) and AFB1+Utt-B. Tricaprylin alone was used as a vehicle control. After the oral administration, the pups were immediately returned to the cage and weaned. Animals were monitored on a daily basis for mortality. Second dose of 6 µg/g body weight of AFB1 was given once the mice were 6 months old. Both males and females were routinely monitored for developmental abnormalities and changes in weight throughout the 12-month study. Drug treatment was started after 5 months after the second dosage of carcinogen. 10mg/Kg Utt-B was administrated orally to the test group on a daily basis continuously for 1 month. After this, animals were sacrificed, and their livers were harvested for histopathology analysis. The livers were examined macroscopically for tumors, and the organs were collected for further studies.

### Isolation of Primary liver cancer cells from AFB1-induced mice liver tumor

Primary liver cancer cells were isolated from the liver tumors developed in DBA/2J mice from the AFB1-induced liver carcinogenesis experiment. Liver tissues were perfused slowly via the portal vein with 20mL of warm Solution1 (HBSS without Ca^2+^ Mg^2+^ and, 0.5 mM EDTA), followed by 20mL of warm Solution 2 (HBSS with Ca^2+^ Mg^2+^, 10 mM HEPES (pH7.4)), followed by 50mL of warm Solution 3 (Solution 2 plus 0.75 mg/ml collagenase D). Following complete digestion of the liver (approx. 15–20 min), the gall bladder was removed and the liver was carefully excised. The tissue samples for isolation of primary cells were collected in ice-cold Hanks Balanced Salt Solution (HBSS) containing antibiotics and processed immediately under sterile condition. Cells were collected from the liver by disrupting the liver capsule and swirling the tissue in a petri dish containing Solution 3. Liver non parenchymal cells (LNPCs) and Kupffer cells were separated from the liver cancer cells by centrifuging the cell suspension at 1000 rpm for 2 min at room temperature. Liver cancer cells in the pellet from the initial centrifugation step were washed in DMEM containing 10% Fetal Bovine Serum (FBS), 200 units/mL penicillin, 200µg/mL streptomycin and 100µg/mL amphotericin B and centrifuged at 1000 rpm for 2 min at room temperature. This washing step was repeated until the supernatant was no longer cloudy [20]. After washing, the cell suspension was transferred to collagen pre-treated 25mm^2^ tissue culture flasks containing DMEM supplemented with 20% FBS, 100 units/mL penicillin and 100µg/mL streptomycin. The cells were grown in a humidified incubator at 37°C with 5% CO_2_ with periodic medium changes. Fibroblast contamination was removed by controlled trypsinization.

### MTT assay

MTT assay was performed in primary liver cancer cells as previously described [21].

### ROS assay

ROS levels within the cells in response to Utt-B treatment were determined by staining the cells using Abcam Cellular ROS kit (ab113851) according to the manufacturer’s protocol.

### Whole transcriptomic sequencing

The Whole Transcriptome Sequencing was performed through the service provider Redcliffe Lifetech Pvt Ltd, Noida, India. RNA was extracted using RNA easy Mini Kit following the manufacturer’s protocol. The RNA quality assessment was done using the RNA ScreenTape System in a 4150 Tape Station System. 1 lL RNA sample was mixed with 3 lL of RNA ScreenTape Sample buffer, then heat denatured at 72 C for 3 min, followed by immediate placement on ice for 2 min. The sample was then loaded onto the Agilent 4150 Tape Station instrument to determine the integrity of RNA using the RNA integrity number (RIN) assigned by the software. The sequence data were generated using Illumina NovaSeq 6000 using fastp v0.20; sequence reads were processed to remove adapter sequences and low quality bases. The QC passed reads were mapped onto the Mus musculus genome using STAR v2 aligner. Using feature-counts software, gene level expression values were obtained as read counts. Spearman Rank Correlation and Principal Components Analysis were used to check whether the biological replicates are corroborating with each other with respect to their expression profiles. Spearman Rank correlation is used for Gaussian distribution, such as gene expression data which shows negative binomial distribution. The biological replicates were grouped as Reference and Test for differential expression analysis. Differential gene expression analysis was carried out using DESeq2 package after normalizing the data using the relative log expression normalization method. Genes with absolute log 2 and fold change above or equal to 1 and adjusted p-value below 0.05 were considered significant. The UpSetR R package was used to generate plots showing overlapping significant genes between conditions. The heat map was generated by using normalized expression values for each sample for a given gene. Gene ontology over representation analysis for biological process was performed using Cluster Profiler R Bioconductor package. Gene Ontology (GO) terms with multiple test adjusted p-value below 0.05 were considered significant.

### Western blotting

Western blotting was performed in tissues and cultured primary liver cancer cells as previously described [21].

### DNA Isolation from tissues

DNA was isolated from both tissues and cultured primary liver cancer cells using DNA Isolation kit for cells and tissues according to manufacturer’s protocol.

### DNA Adduct analysis

DNA adduct formation in control and Utt-B treated samples (both tissues and primary liver cancer cells) was evaluated using Cell Biolabs’ Aflatoxin DNA Adduct Competitive ELISA Kit, according to manufacturer’s protocol.

### RNA isolation

RNA isolation from cells and tissues were done using NucleoSpin RNA, Mini kit for RNA purification from Macherey-Nagel according to manufacturer’s protocol.

### First strand synthesis

First strand synthesis was carried out using High-capacity cDNA synthesis kit according to the manufacturer’s protocol. 2μg of RNA was used for the synthesis.

### Real-Time PCR

Real-Time PCR was performed using Power SYBR Green PCR master mix, Applied Biosystems. The housekeeping gene, GAPDH, was used as reference gene for normalization. The data were calculated using the 2^-ΔΔCt^ method [22].

### Preparation of S9 fractions

The rat livers were collected from Wistar rats as per protocol (IAEC/717/RUBY/2018) approved by the Institutional Animal Ethics Committee, Rajiv Gandhi Centre for Biotechnology. Rat liver microsomal fraction was prepared according to the Garner method [23].

### Ames test

Ames test was performed as previously described [24]. The inhibition of mutagenesis was expressed as the percentage of inhibition of mutagenic activity

### Histology and Immunohistochemistry

Haematoxylin and eosin staining was performed as previously described. Immunohistochemical analysis was done using Poly Excel HRP/DAB detection system universal kit as per manufacturer’s protocol [16].

### Sirius Red Staining

Sirius red staining was performed as previously described [16].

### Immunofluorescence

Immunofluorescence analysis was performed as previously described [15].

### Statistical analysis

Data represent the results for assays performed in triplicate. H-scoring for Immunohistochemistry and the quantification of western blot were carried out using ImageJ software. The statistical analysis was performed using Graph Pad Prism software (Graph Pad Software Inc., San Diego, CA, USA). The error bars represent ± SD, taken from the three independent experiments.

## Results

### Uttroside B inhibits the development of hepatic tumors in DBA/2J mice from aflatoxin B1-induced liver carcinogenesis model

Previous studies conducted in our lab have demonstrated the chemotherapeutic potential of Utt-B against HCC using sub-cutaneous and orthotopic xenograft models established in immuno-suppressed mice [14–16, 25]. However, these models do not provide a comprehensive understanding of the influence of Utt-B on the immune landscape of HCC. Hence, in the present study, we established an AFB1-induced environmental liver carcinogenesis model in immuno-competent DBA/2J mice to evaluate the immunomodulatory effects of Utt-B against HCC [**Figure 1 A**]. The animals from the positive control group exposed to AFB1-alone showed severe weakness, and four animals died towards the end of the study period. However, no mortalities were observed in the Utt-B treatment group. While the animals in the negative control group that were not exposed to either AFB1 or Utt-B did not display any signs of tumor development throughout the study period, all the animals from the AFB1 group developed solid hepatic tumors. None of the animals in the Utt-B-treated group developed solid tumors [**Figure 1 B-C**]. Notably, the body weight of animals in AFB1-alone control group were lower as compared to that of Utt-B-treated animals [**Figure 1 D**]. The average liver weight and the liver: body weight ratio was significantly decreased in the animals from AFB1-alone group as compared to the negative control group possibly due to the AFB1-induced liver injury. However, the average liver weight and the liver: body weight ratio in the Utt-B treated group, was comparable to the negative control group [**Figure 1 E-F**]. The organs excised from different experimental group showed splenomegaly in some of the AFB1-exposed mice [**Supplementary Figure 1A**]. Additionally, the elevated levels of AFP and total cholesterol were stabilized in AFB1-exposed mice following Utt-B treatment [**Figure 1 G-H**]. Serum biochemical analysis revealed high levels of liver enzymes such as AST, ALT, and ALP in AFB1-administered animals. Utt-B treatment significantly reduced this abnormally increased liver enzyme activity [**Figure 1 I**]. Furthermore, Utt-B treatment lowered the elevated levels of urea, uric acid, and creatinine that was observed in AFB1-exposed mice [**Figure 1J**]. Histopathological assessment of liver tissues revealed the presence of regenerative hyperplasia and atypical nodules in AFB1-exposed animals. Dysplastic hepatocytes that are indicative of carcinoma, were more prominent in the livers of AFB1-exposed mice, while the livers from mice treated with Utt-B showed normal histology [**Figure 1K**]. Additionally, the presence of focal lymphoid aggregates was noted in the lung tissues of AFB1-exposed animals that was absent in mice treated with Utt-B [**Supplementary Figure 1B**].

**Figure 1:**
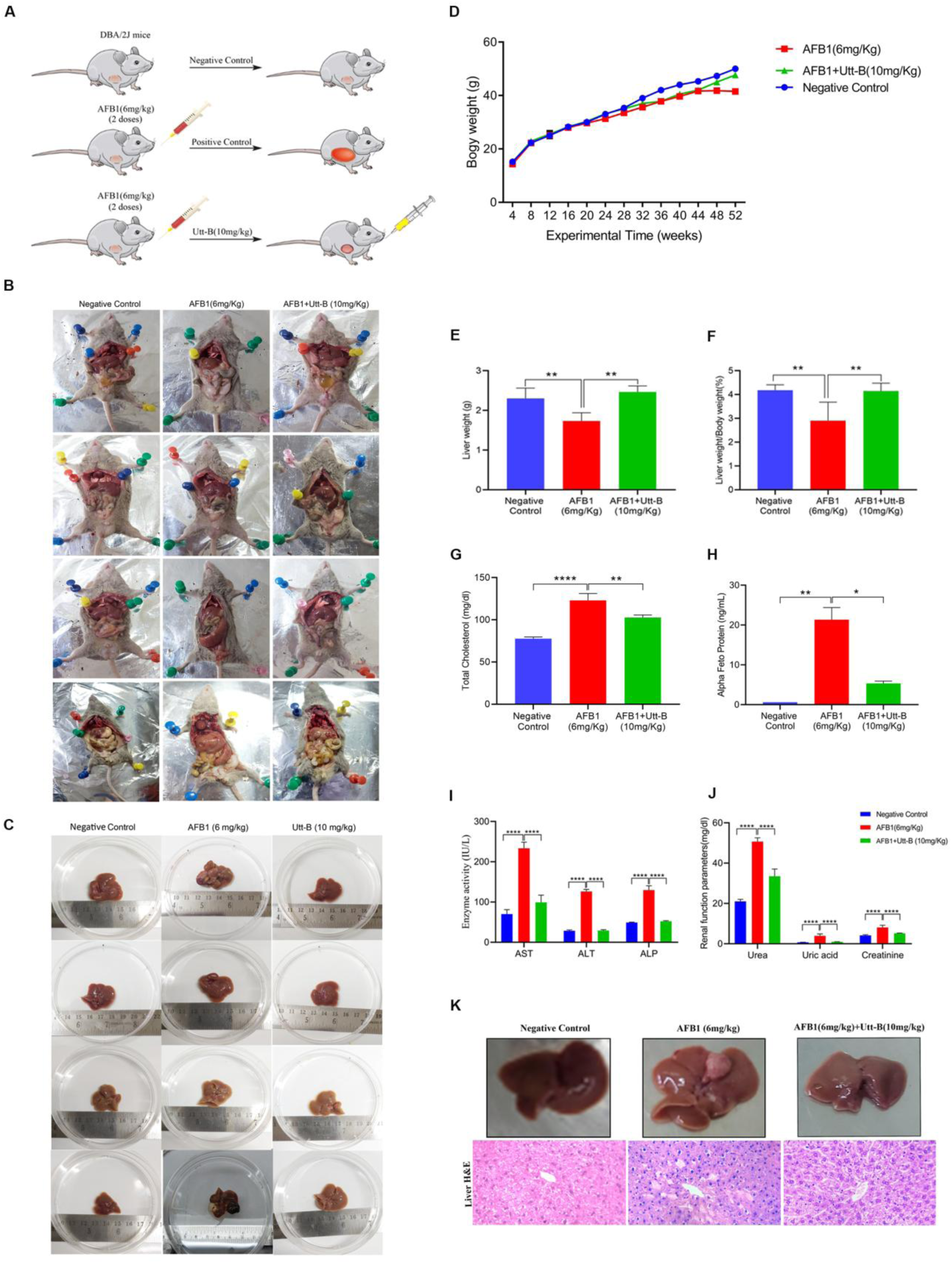
Utt-B inhibits the development of hepatic tumors in mice exposed to AFB1. **(A)** The schematic diagram representing the experimental design of AFB1-induced liver carcinogenesis in DBA/2J mice **(B)** Representative photographs of mice from different groups of AFB1-induced liver carcinogenesis model **(C)** Representative photographs showing the gross morphology of tumor-bearing liver tissues from different groups of AFB1-induced liver carcinogenesis model **(D)** Graph showing average weekly body weight (g) of animals in different treatment groups post-AFB1 administration. **(E, F)** The average weight of liver (g) and liver weigh:body weight (%) in Control, AFB1-alone, and AFB1+Utt-B groups. The error bars represent ±S.D. Data shows the average of three independent sets of experiments with 6 animals per group. One-way ANOVA followed by Tukey’s test was used for statistical comparison between different groups **p≤0.01. **(G, H)** The serum level of total cholesterol and AFP in different groups of mice treated with AFB1-alone and AFB1+Utt-B and normal DBA/2J mice. The graphs represent mean± S.D. One-way ANOVA followed by Tukey’s test was used for statistical comparison between different groups **p≤0.01. **(I,J)** The serum level of liver function enzymes and renal function parameters of mice from different groups. The graphs represent mean±S.D. One-way ANOVA followed by Tukey’s test was used for statistical comparison between different groups ****p≤0.0001. **(K)** Histopathological changes induced by AFB1 administration in liver tissues isolated from different treatment groups of AFB1-induced liver carcinogenesis model.

### Treatment with uttroside B significantly attenuates aflatoxin B1-induced liver injury and cancer progression in a murine model of hepatocarcinogenesis

Collagen I is a vital component of the liver extracellular matrix and its excessive accumulation in the liver tissues are indicative of tumor growth, angiogenesis, and metastatic progression of HCC [26, 27]. Collagen I expression is up-regulated in 83.7% of human HCC specimens compared to adjacent non-tumor tissues [27, 28]. Notably, our results indicate that Utt-B significantly impedes collagen accumulation in the liver tissues as compared to AFB1-alone group as evident from Sirius Red staining that stains collagen I and III fibres. Further, Utt-B-treated liver tissues showed a significant decrease in the mRNA expression levels of Collagen 1-α as compared to liver tissues of mice from AFB1-alone group [**Figure 2 A**]. The relative expression of SREBP-1c, SCD-1 and MCP-1, some of the key genes associated with liver injury and progression to HCC were found to be significantly down-regulated in Utt-B-treated tissues in comparison with AFB1-alone tissues. The list of all primer sequences used for qRT-PCR analysis have been enlisted in **Supplementary table 1-2**. Further, the analysis of proliferative biomarker, PCNA, and the HCC specific biomarker, AFP, revealed a significant down-regulation of these biomarkers in the Utt-B-treated liver tissues as compared to AFB1-alone group [**Figure 2 C-E**]. AFB1-induced HCC is highly associated p53 gene mutation. Immunohistochemical analysis showed a substantial increase in the tissue expression levels of the tumor suppressor genes, p53, and its downstream target, p21, in Utt-B treated liver tissues as compared to AFB1-exposed group [**Figure 2 F**]. Further, an 8-fold increase in the relative gene expression of p53 and 5.4-fold increase in the expression of p21 was observed in the liver tissues of Utt-B-treated mice from the AFB1-induced liver carcinogenesis model [**Figure 2 G**].

**Figure 2:**
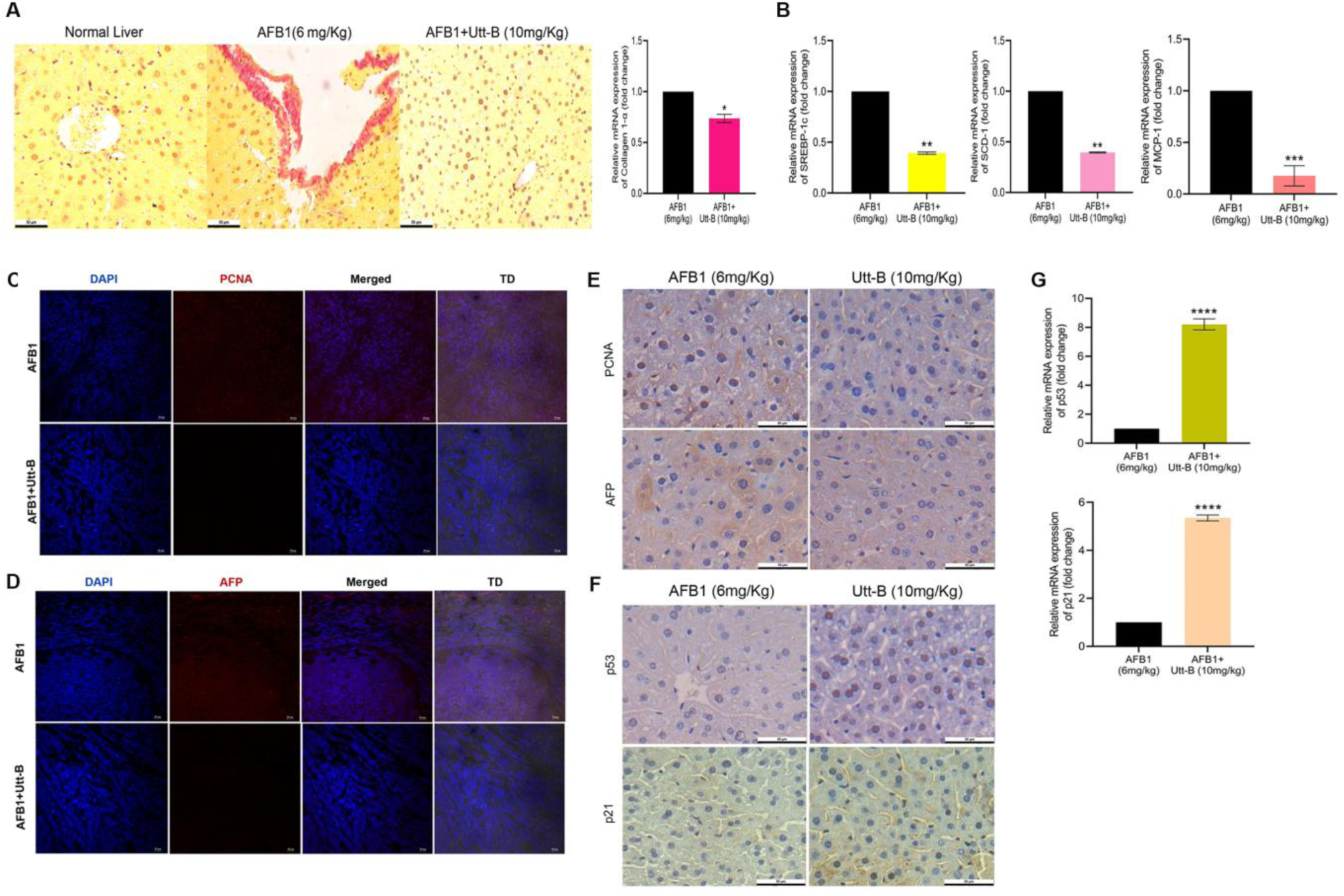
Utt-B attenuates AFB1-induced liver injury and cancer progression in mice. **(A)** Utt-B significantly inhibits the collagen deposition in liver tissue surrounding the portal vein as evident from Sirius red staining and down-regulation of Collagen-1a mRNA expression levels in the tissue **(B)** Utt-B down-regulates the expression of SREBP-1c, SCD-1 and MCP-1 genes associated with HCC pathogenesis **(C-E)** Utt-B significantly down-regulates PCNA and AFP in the liver tissues of mice from the AFB1-induced HCC model. **(F-G)** Utt-B significantly up-regulates p53 and p21 in the liver tissues of mice from the AFB1-induced HCC model. The graphs represent mean±S.D. One-way ANOVA followed by Tukey’s test was used for statistical comparison between different groups. *P ≤ 0.05, **P ≤ 0.01,***P ≤0.001, ****P ≤ 0.0001.

### Uttroside B is cytotoxic towards primary liver cancer cells isolated from the hepatic tumors of mice exposed to aflatoxin B1

Primary liver cancer cells were isolated from the AFB1-induced hepatic tumors from DBA/2J mice as described in “Materials and methods” [**Figure 3A-B**]. Immunofluorescence analysis of proliferative markers such as Ki-67, PCNA, the epithelial cancer cell marker, pan-cytokeratin and the HCC specific biomarker AFP, revealed high expression in AFB1-derived primary cancer cells, indicating the rapid proliferative nature of these cells and validating the epithelial origin of these primary cancer cells [**Figure 3 C-F**]. Further, we checked the dose-dependent cytotoxicity of Utt-B and the FDA-approved anti-HCC drug, sorafenib towards primary liver cancer cells using MTT assay. Although, both Utt-B and sorafenib were found to be cytotoxic towards the primary cancer cells, Utt-B with an IC50 value of 4µM was found to be 3-fold more cytotoxic towards the primary liver cancer cells than sorafenib which had an IC50 value of 12µM [**Figure 3 G-H**]. Furthermore, the relative mRNA expression levels of p53 and p21 were significantly up-regulated in the AFB1-derived primary cancer cells treated with Utt-B, thereby validating the consistent tumor suppressive effects of the compound in the primary liver cancer cells [**Figure 3 I**].

**Figure 3:**
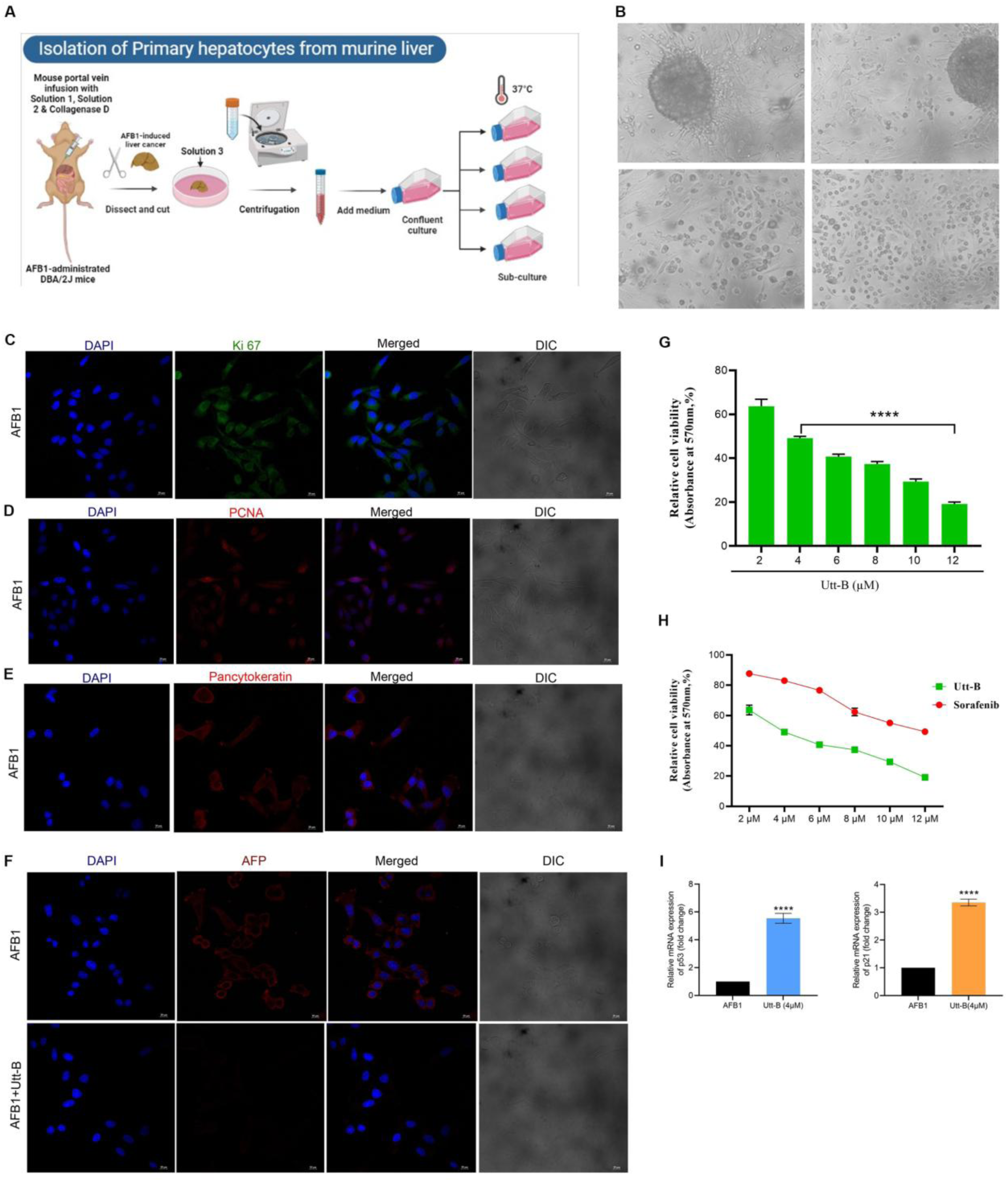
Utt-B exhibits cytotoxicity towards primary cancer cells isolated from the livers of mice from AFB1-induced liver carcinogenesis model. **(A)** Schematic diagram showing the isolation of primary cells from AFB1-induced liver cancer model **(B)** Phase contrast images of primary liver cancer cells isolated from AFB1-induced liver tumor in DBA/2J mice **(C-E)** Primary liver cancer cells isolated from AFB1-induced hepatic tumor showing increased expression of Ki-67, PCNA and pan-cytokeratin **(F)** Utt-B significantly down-regulates the expression of AFP in primary liver cancer cells isolated from AFB1-induced hepatic tumor. **(G)** Utt-B is cytotoxic towards primary cells isolated from AFB1-induced liver tumors **(H)** Primary cells isolated from AFB1-induced liver tumors were treated with varying concentrations of Utt-B and sorafenib and the cell viability was assessed by MTT assay **(I)** Utt-B treatment up-regulates the expression of p53 and p21 in primary liver cancer cells. The error bars represent ± S.D. One-way ANOVA followed by Tukey’s test was used for statistical comparison between different groups. ****P ≤ 0.0001.

### Uttroside B significantly inhibits the AFBO-DNA Adduct formation in aflatoxin B1-induced liver carcinogenesis model

Aflatoxin B1 is a genotoxic hepatocarcinogen, that causes cancer by inducing DNA adduct formation, which lead to genetic changes in target liver cells. Aflatoxin-8,9-epoxide (AFBO), a reactive metabolite formed after AFB1 metabolism by cytochrome P450 is capable of intercalating into DNA at the N7 position of guanine. As a result, the guanine is released from the DNA, leaving an apurinic site, and the ribose ring opens forming a stable formamido pyrimidine (FAPY) adduct. Previous literature has documented the presence of the FAPY adduct in the livers of rat which have been subjected to AFB1 exposure. Previous studies using animal models have also demonstrated a linear dose-response relationship between AFB1-DNA adduct formation and development of liver tumors. AFB1 exposure is known to induce mutation in the codon 249 in the p53 tumor suppressor gene wherein the AGG (arginine) residue is changed to AGT (serine) residue. This mutation results in the inactivation of p53 protein [**Figure 4 A**]. DNA adduct analysis was performed in the liver tissues from AFB1-induced liver carcinogenesis model as well as from primary cells isolated from AFB1-induced liver tumors of DBA/2J mice. Interestingly, Utt-B treatment resulted in a significant reduction in the concentration of AFBO-DNA adducts in the livers of DBA/2J mice as well as in the primary cells isolated from the liver tissues of these mice, thereby clearly indicating the chemotherapeutic potential of Utt-B in AFB1-induced liver carcinogenesis. The concentration of AFBO-DNA adducts in the tissue samples in AFB1-alone group was 4.170 µM, while it was only 1.12 µM in Utt-B treated group [**Figure 4 B**]. Similarly, the concentration of AFBO-DNA adduct in the control primary cells was 3.16µM and it reduced to 1.5 µM in the Utt-B treated group [**Figure 4 C-D**].

**Figure 4:**
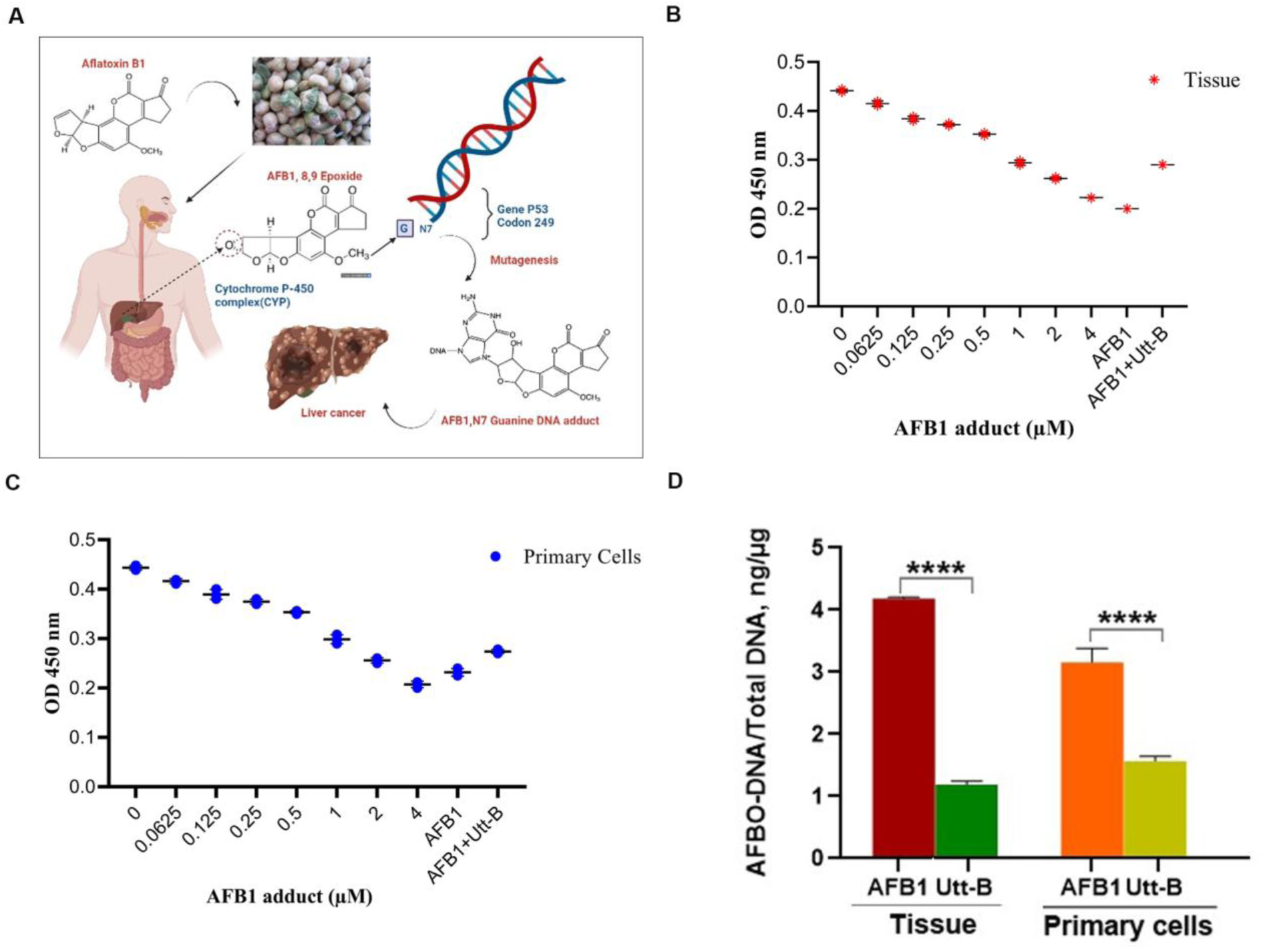
Utt-B mitigates AFB1-induced DNA adduct formation, both *in vitro* and *in vivo*. **(A)** Schematic diagram showing the AFBO-adduct formation in AFB1-induced liver carcinogenesis. **(B)** Utt-B treatment significantly reduces the concentration of AFBO-DNA adducts formed in the liver tissue of DBA/2J mice treated with Utt-B. Standard curve represents the AFBO-DNA adduct formation in liver tissues isolated from different treatment groups of AFB1-induced liver carcinogenesis model **(C)** Utt-B treatment significantly reduces the concentration of AFBO-DNA adducts formed in the primary cells isolated from liver tissue of DBA/2J mice, treated with AFB1. **(D)** Comparison of the efficacy of Utt-B against AFBO-DNA adduct formation in liver tissues and primary liver cancer cells. The error bars represent±S.D. One-way ANOVA followed by Tukey’s test was used for statistical comparison between different groups. ****P ≤ 0.0001.

### Uttroside B potentiates immunogenic apoptosis against hepatocellular carcinoma

Our next attempt was to evaluate the immunomodulatory effects of Utt-B against HCC. *In vitro* studies conducted using HepG2 cells indicate that Utt-B induces the production of ROS starting from 30 min post-treatment and is sustained until 24 h post-treatment [**Figure 5A**]. Interestingly, a strong-up-regulation of HSP70 was observed in the HepG2 cells treated with Utt-B [**Figure 5B**]. Furthermore, it was observed that Utt-B caused a marked increase in the mRNA expression of the inflammatory cytokine IL-1β and the chemokine CXCL10 in HepG2 cells [**Figure 5C**]. Collectively, the current *in vitro* findings highlight Utt-B as a potential inducer of immunogenic cell death against liver cancer cells. Interestingly, a significant increase in the relative mRNA expression levels of Annexin A1, an ICD-related DAMP, was observed in the tissues of Utt-B treated mice in comparison to the control group from the AFB1-induced liver carcinogenesis model. This observation suggests the potential of Utt-B in triggering immunogenic cell death, *in vivo* [**Figure 5 D**]. Whole transcriptome sequencing analysis was performed using the liver tissues in order to delineate the immunological changes induced by Utt-B in the TME. A total of 37 genes were found to be down-regulated and 38 genes were up-regulated in the Utt-B treatment group compared to the control group [**Figure 5 E**]. Further, the functional gene set enrichment analysis of the transcriptomics data set was performed using Gene ontology in iDEP2.0 web application. An up-regulation in the positive regulation of dendritic cell-mediated antigen processing and presentation was noted in the liver tissues of mice treated with Utt-B. Further, an up-regulation in the MHC-II protein complex assembly and peptide antigen assembly with MHC-II complex was also observed in the Utt-B treated tissues [**Table 1**]. The enhancement in the MHC complex assembly and enhancement in the antigen presentation in the TME upon Utt-B treatment was further confirmed using the qRT-PCR analysis of the expression levels of B2M, PSMB8, and LGMN. B2M is a protein that forms a part of MHC-I molecules and PSMB8 is involved in the processing of tumor antigen peptides for being presented by MHC-I molecules [29]. A significant increase in the transcription of these molecules was noted upon treatment with Utt-B. Likewise, it was also noted that Utt-B induces the transcriptional activation of LGMN and CD74, key players in mediating the processing of endogenous antigens and presentation of antigens by MHC-II [**Figure 5 F**]. Hence, the current results validate the role of Utt-B in improving antigen processing and presentation to T cells in the TME of HCC. Interestingly, a drastic enhancement in leukotriene B4 (LTB4) metabolism was observed in Utt-B-treated group. Activated DCs (Dendritic Cells) are known to induce the chemotactic attractant, LTB4, which in turn activates transcription of cytokine genes. LTB4 serves as a bridge that connects innate and adaptive immunity by promoting infiltration of cytotoxic T lymphocytes (CTLs), which in turn promote immune surveillance leading to cytotoxicity and death of tumor cells [30]. Chemotherapeutic agents inducing ICD have been reported to induce ROS production, autophagy and endoplasmic reticulum stress. It is well known that the unfolded protein response switches from a cell survival to a cell death strategy when the ER homeostasis is lost [31]. Notably, our functional Gene set enrichment analysis of the transcriptomics data also revealed that Utt-B stimulates ER stress and unfolded protein response in the TME [**Table 1**]. Therefore, the occurrence of Utt-B-mediated ER stress and ROS production could be one of the underlying mechanisms that contribute to immune-activation and immunogenic apoptosis induced by the compound in the context of HCC.

**Figure 5:**
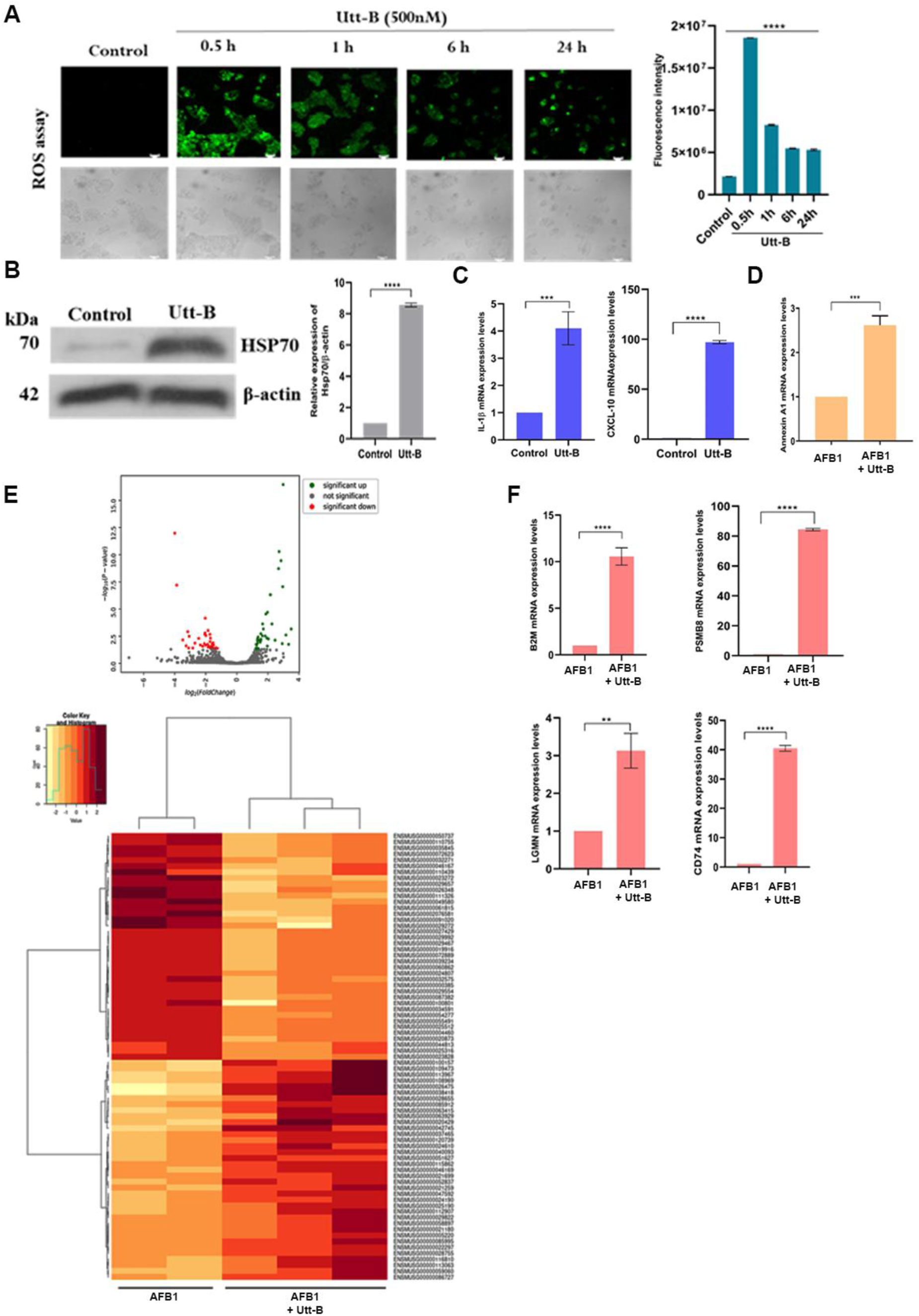
Utt-B induces the production of DAMPs and enhances antigen presentation in the TME of AFB1-induced liver cancer. **(A)** Treatment with Utt-B leads to the production of reactive oxygen species in HepG2 cells **(B)** Utt-B elevates the expression of HSP70 in HepG2 cells **(C)** Utt-B up-regulates the expression of IL-1β and CXCL10 in HepG2 cells **(D)** Utt-B up-regulates the expression of ANXA1 in the tumor tissues from AFB1-induced liver carcinogenesis model **(E)** Volcano plot and heatmap showing differential gene expression between AFB1-alone and Utt-B treated tumor tissues from the AFB1-induced liver carcinogenesis model using Whole transcriptomic sequencing **(F)** Utt-B up-regulates the expression of B2M, PSMB8, LGMN and CD74 genes in the tumor tissues from AFB1-induced liver carcinogenesis model. The error bars represent ± S.D. One-way ANOVA was used for statistical comparison between different groups. ****P ≤0.0001; ***P ≤ 0.001; **P ≤ 0.01.

**Table 1:**
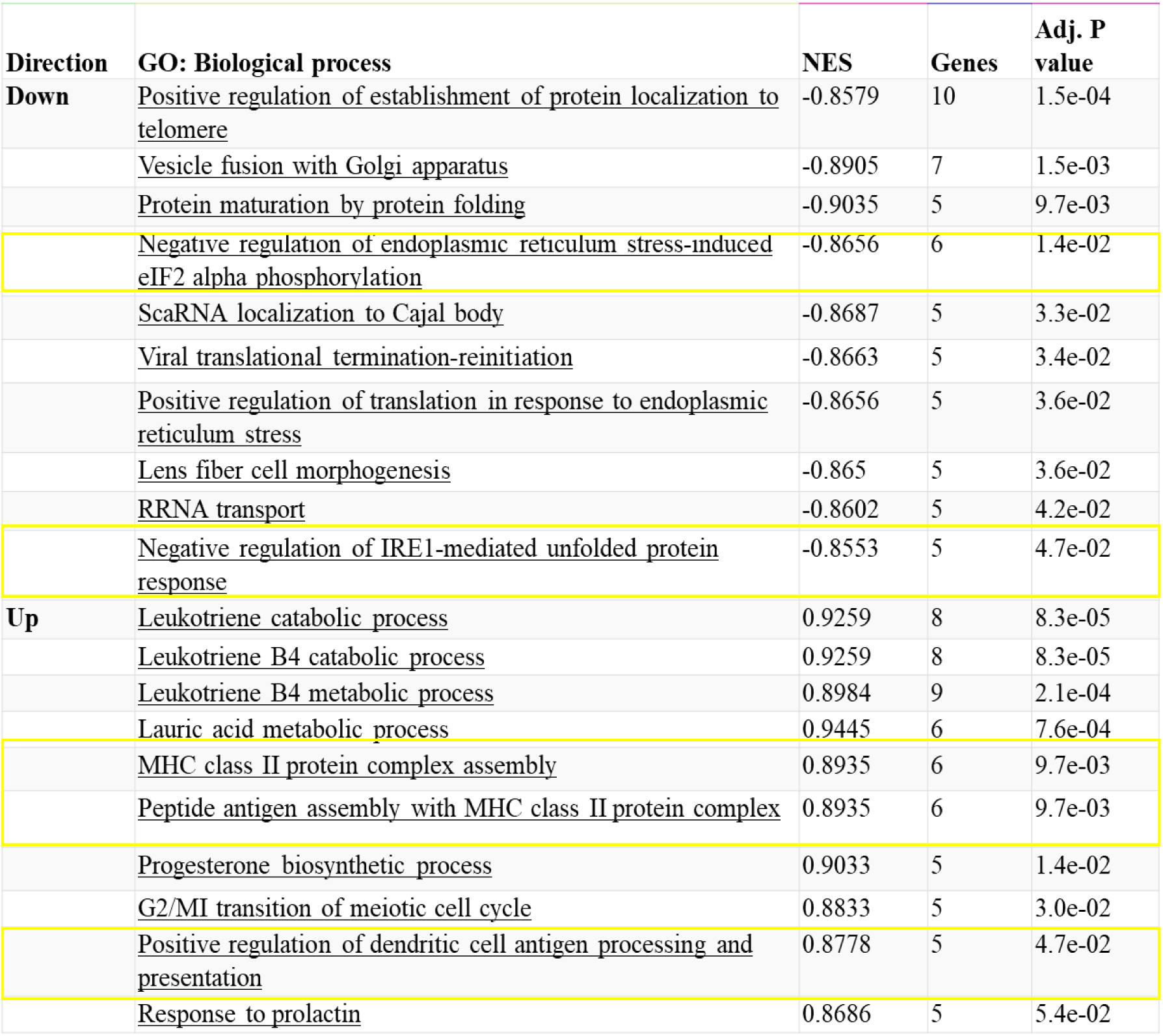
GSEA Pathway analysis of RNA Sequencing data depicting the differential regulation of biological processes in the tumor tissues from the AFB1-induced liver carcinogenesis model in response to Utt-B treatment.

Additionally, the changes in the expression of T_h_1, T_h_2 and T_h_17 cytokines in the liver tissues were also evaluated using qRT-PCR analysis. An increase in the levels of T_h_1 cytokines aids in the activation of cytotoxic T cells, whereas the increase in the T_h_2 and T_h_17 cytokines favor the activation of T helper cells. From the clinical perspective, skewing of the TME towards the T_h_1 phenotype is associated with good prognosis in HCC patients. The present results reveal that treatment with Utt-B increases the transcriptional activation of T_h_1 cytokines, namely IL-2, TNF-α and IFN-γ. A significant reduction in the levels of T_h_2 cytokines, IL-4, IL-6 and IL-10 and T_h_17 cytokine, IL-17A in response to Utt-B treatment was also observed [**Figure 6 A**]. These results suggest that Utt-B shifts the TME towards T_h_1 phenotype, associated with good prognosis. The positive interaction between the professional antigen presenting cells (APCs) such as DCs and T cells is dampened by the negative interaction of immune checkpoint molecules present on the surface of T cells with their ligands on the tumor cells. This negative interaction sustains the T cells in an exhausted state, thereby inhibiting anti-tumor effector responses in the TME. Hence, we evaluated whether Utt-B is instrumental in impeding this negative interaction and it was observed that Utt-B significantly reduces the transcriptional activation of the major immune checkpoints and T-cell exhaustion markers, namely, CTLA-4, PD-1, TIM-3, LAG-3 and TOX in the liver tumors [**Figure 6 B**]. Evidently, treatment with the compound attenuates T cell exhaustion, and paves the way for induction of anti-tumor effector responses*, in vivo*. DCs are known to stimulate specific T-cell mediated anti-tumor response owing to their ability to cross present tumor associated antigens to naïve T cells. The recognition of antigen by the T cell receptor triggers an array of downstream signalling cascades, which are instrumental in eliciting anti-tumor effector T cell responses. ZAP70 and Lck are the major kinases that activate T cell signalling downstream of TCR. Interestingly, it was observed that Utt-B induces the transcriptional activation of both these kinases in the tumor tissues, suggesting the potential role of Utt-B in triggering an anti-tumor effector T cell response [**Figure 6 C**]. CTL-mediated apoptosis occurs primarily via activation of the serine protease, granzyme B. The activated granzyme B in turn, contributes to the induction of apoptosis. Immunohistochemical analysis of granzyme B was performed in the tumor tissues from AFB1-induced carcinogenesis model and the results demonstrate the elevated expression of granzyme B in the Utt-B treated tumor tissues as compared to the control tumor tissues [**Figure 6 D**]. Utt-B treatment triggers apoptosis, as evident from the cleavage of caspase 9 [**Figure 6 E**] and PARP [**Figure 6 F**] in liver tissues, in contrast to the AFB1-alone control group. Furthermore, the elevated expression of cleaved PARP in the Utt-B treated group confirms the occurrence of apoptosis in the liver tissues, thereby, validating the anti-tumor immunomodulatory effects of Utt-B in the TME of HCC [**Figure 6 G**]. Taken together, the present results demonstrate the complementary action of the immunogenic cell death induced by Utt-B in augmenting the chemotherapeutic, anti-carcinogenic and pro-apoptotic effects of the compound in the context of HCC.

**Figure 6:**
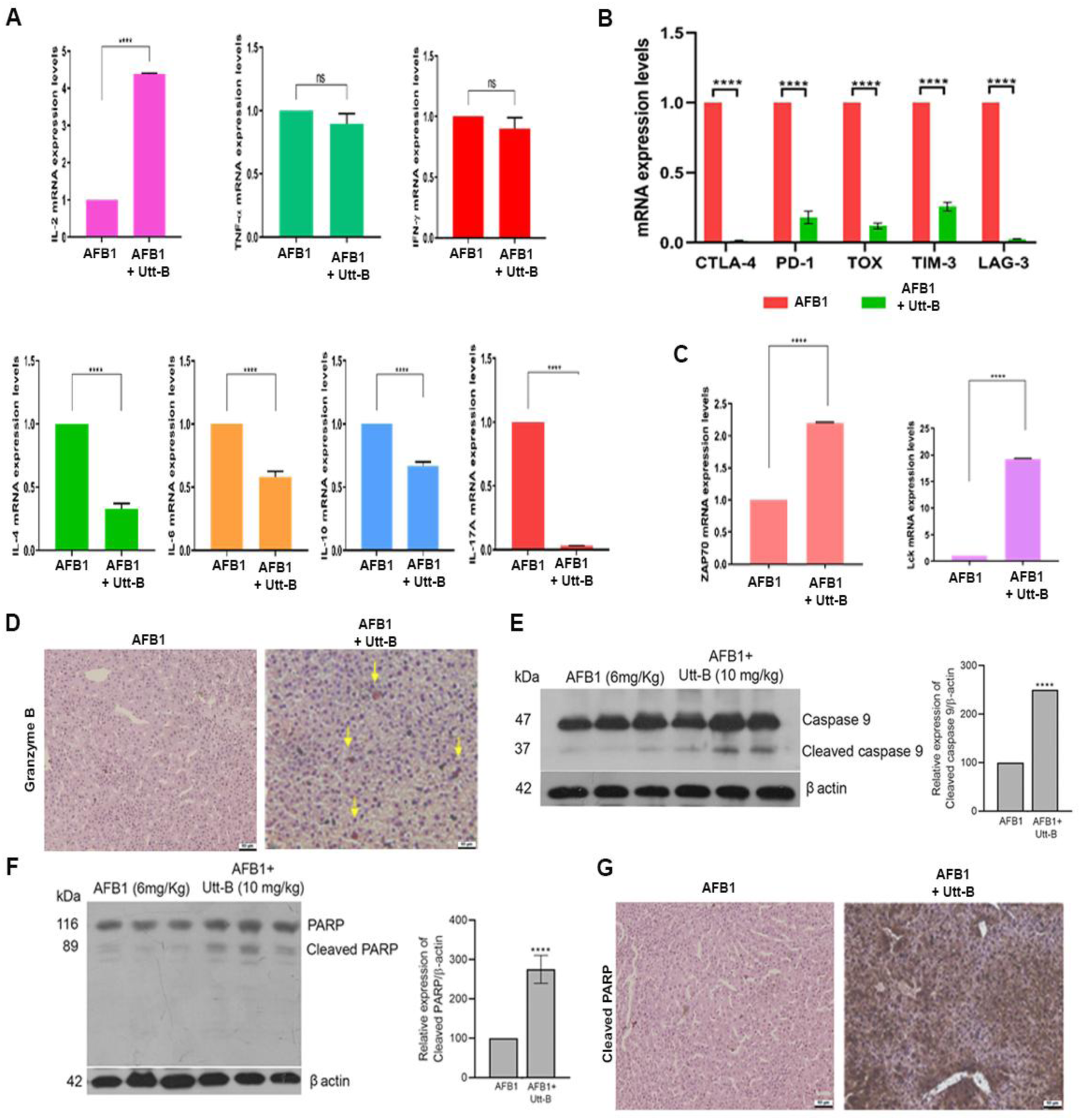
Utt-B potentiates immunogenic apoptosis in the livers of mice from the AFB1-induced liver carcinogenesis model. **(A)** Utt-B skews the TME towards the Th1 phenotype **(B)** Utt-B down-regulates the mRNA the expression of major immune checkpoints **(C)** Utt-B up-regulates the mRNA expression of major T cell activation kinases, ZAP70 and Lck **(D)** Utt-B significantly up-regulates granzyme B in the liver tissues of mice from the AFB1-induced HCC model **(E-G)** Utt-B significantly induces the cleavage of caspase 9 and PARP in the liver tissues of mice from the AFB1-induced liver carcinogenesis model. The error bars represent ± S.D. One-way ANOVA was used for statistical comparison between different groups. ****P ≤ 0.0001; ***P ≤ 0.001; **P ≤ 0.01.

### Uttroside B significantly inhibits aflatoxin B1-induced mutagenesis of TA98 strain of *S. typhimurium*

Metabolism of parent molecules in the liver is often associated with the production of mutagenic metabolites as the by-products of metabolism in the liver. In the present study rat liver S9 fraction is used to mimic the mammalian metabolic conditions for assessing the mutagenic potential of metabolites formed by a parent molecule in the hepatic system. The TA98 strain of *S. typhimurium* bacteria contain a mutation at histidine operon, which makes them incapable of synthesizing enzymes vital for the biosynthesis of histidine. Therefore, these strains are incapable of growing in a histidine deficient environment. When these strains are exposed to a mutagen, they undergo genetic reversion that capacitates their growth in histidine deficient environment. Thus, the mutagenicity of a compound can be assessed by the capability of increasing the clonogenic potential of *S. typhimurium* in medium that lacks histidine. Our current results suggest the anti-carcinogenic potential of Utt-B. Since AFB1 is a potent mutagen known to induce DNA damage and p53 mutation, we evaluated whether Utt-B is capable of preventing AFB1-induced mutagenesis, contributing to its efficacy to prevent AFB1-induced liver carcinogenesis. TA-98 strain of *Salmonella typhimurium* was used for the Ames test since it is known to be sensitive towards aflatoxin B1 and contains a mutagenesis enhancing plasmid, pKM101. Previous literature suggests that mutagenesis of AFB1 is detectable at a range of 10-50ng. An increase in the concentration above 100ng, leads to the killing of the tester strains and in turn results in the decrease in the PFU/mL [32, 33]. Based on the previous reports, 50ng of AFB1 was chosen for the current study. 2-Acetamidofluorine (30µg/plate) was used as the positive control. Different concentrations of Utt-B i.e., 5µg, 10 µg, and 25µg were used for assessing the potency of Utt-B in blocking AFB1-induced reversion. Utt-B was incubated with the S9-fraction containing TA98 and 0.05µg of AFB1 along with co-factors required for *in vitro* xenobiotic metabolism. Efficacy of Utt-B to revert the mutagenic effect of AFB1 which converts the His2 strains of *S. typhimurium* TA 98 to His1 in the presence of S9, a rat liver fraction containing cytochrome P450, was analysed. Although a significant increase of mutagenesis was observed in the AFB1-treated strains of *S. typhimurium,* Utt-B could inhibit the mutagenesis induced by AFB1 in a dose-dependent manner with significant reduction at a concentration of 25µg/plate [**Figure 7 A**]; [**Supplementary Table 3**]. The inhibition of mutagenesis expressed as the percentage of inhibition of mutagenic activity is depicted in **Figure 7 B**. The findings of the Ames test suggest that Utt-B could have a potential role in preventing the initiation step of HCC.

**Figure 7:**
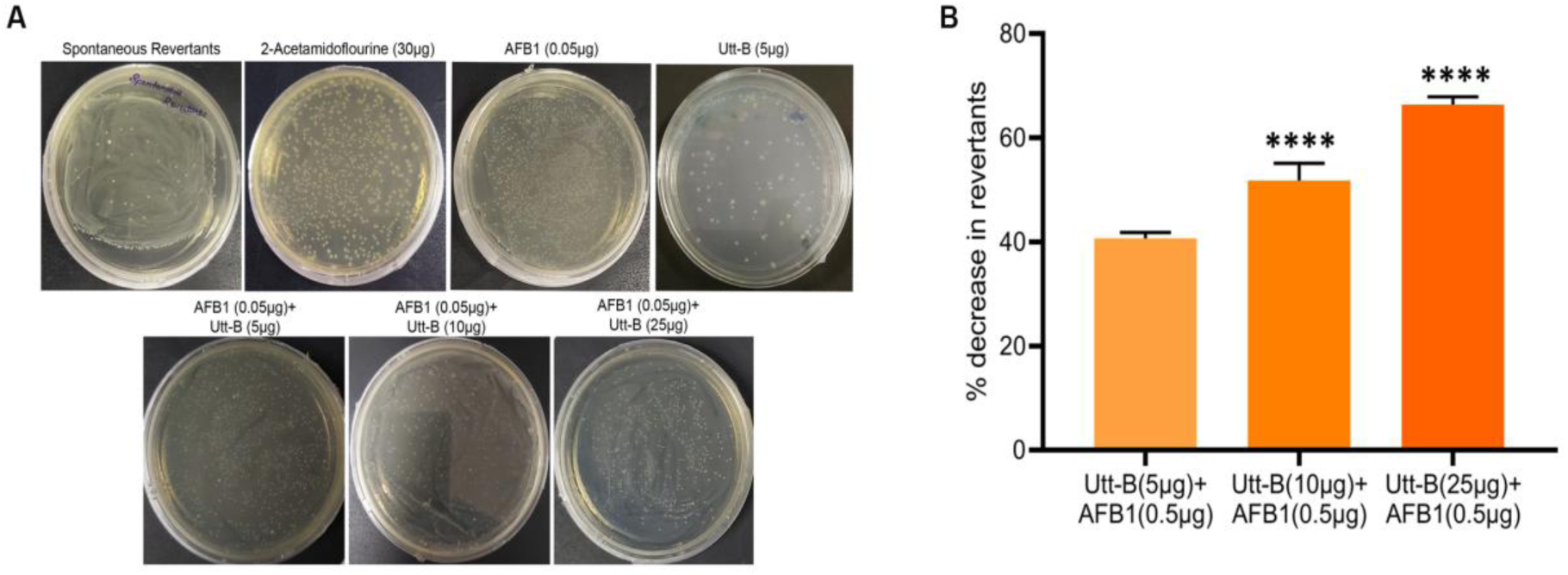
Utt-B inhibits the AFB1-induced mutagenesis of TA98 strain of *S. typhimurium*. **(A)** The anti-mutagenic activity of Utt-B against AFB1 evaluated using Ames test **(B)** Graphical representation of percentage of spontaneous revertants upon increasing concentration of Utt-B. The error bars represent±S.D. One-way ANOVA followed by Tukey’s test was used for statistical comparison between different groups. ****P ≤ 0.0001.

## Discussion

The biotransformation of AFB1 majorly occurs in the liver. This leads to hepatotoxicity and hepatocyte damage [34]. AFB1 exposure in male Kunming mice reduced both absolute and relative liver weights, along with increased levels of liver enzymes AST, ALP, ALT, and AFP [35, 36]. Our results suggest that treatment with Utt-B significantly mitigates elevated levels of liver enzymes. It is well known that aflatoxin can interact with DNA base pairs through its active epoxide moiety, and in turn cause genetic mutations in the p53 gene. Mutant p53 proteins may facilitate selective clonal expansion of hepatocytes during carcinogenesis. Hepatocellular carcinoma often exhibits reduced expression of p21 and p53. p21 is pivotal in p53-mediated cell cycle arrest in response to DNA damage. In a previous study involving HCC patients, 38 out of 59 patients showed altered p53 expression and the absence of p21 protein. Hence, the absence of p21 protein in HCC was associated with altered p53 expression [37]. Our results suggest that Utt-B treatment notably up-regulates the expression of p53 and p21, two major tumor suppressor genes, in both the tumor tissues of mice from AFB1-induced liver carcinogenesis model as well as the primary liver cancer cells isolated from AFB1-induced liver tumors. Previous literature indicates that AFB1 induces DNA adducts, leading to genetic changes in liver cells. AFBO-DNA adducts serve as biomarkers for evaluating AFB1-induced liver damage. A previous study has reported that curcumin alleviates AFB1-induced liver damage in chicken, suggesting the therapeutic effects of phytochemicals [38]. Similarly, treatment with Utt-B led to a marked decrease in AFBO-DNA adducts in DBA/2J mice exposed to AFB1, indicating its ability to ameliorate AFB1-induced liver damage.

AFB1 induces immunosuppression in hosts by diminishing DC-mediated antigen presentation [39]. ICD is a relatively new form of regulated cell death, one that is capable of eliciting an anti-tumor adaptive immune response [40]. Previous literature suggests the ICD-inducing potential of various phytochemicals such as, curcumin, resveratrol, silibinin, capsacin, ginsenoside and plumbagin [41–43]. It is well known that ICD inducers cause extensive reactive oxygen species (ROS) production in cancer cells which is responsible for triggering intrinsic apoptosis, that culminates in ICD [44, 45]. The present results suggest that treatment with Utt-B prevents the development of AFB1-induced hepatic tumors in mice and attenuates the immunosuppression in the TME via induction of ICD. Interestingly, chemotherapeutic agents are known to induce irreparable ER homeostasis, and in turn transform the unfolded protein response (UPR) from a pro-survival to a pro-apoptotic cell death program leading to ICD [31]. The *in vitro* and *in vivo* findings of the current study collectively suggest that Utt-B induces ROS production, and positively regulates UPR. This could be one of the mechanisms by which Utt-B induces ICD against HCC. A recent report by Khedr *et al*., reveals that berberine-loaded albumin nanoparticles reverse aflatoxin B1-induced liver hyperplasia via induction of apoptosis [46]. Interestingly, Utt-B treatment led to the induction of apoptosis in the AFB1-induced liver cancer model, as is evident from the cleavage of caspase 9 and PARP. The current findings demonstrate that apart from its chemotherapeutic potential, Utt-B also possesses a remarkable immunomodulatory function which complements its anti-tumor and pro-apoptotic efficacy against HCC. Notably, the results of the Ames test indicate that Utt-B exhibits anti-mutagenic property by inhibiting the mutagenic potential of AFB1 in the TA98 strain of *S. typhimurium*. Our results indicate that Utt-B counteracts mutations induced by AFB1 in a concentration-dependent manner, marking its anti-mutagenic property and anti-cancer activity in an environmental carcinogenesis model. However, further studies involving pre-clinical chemical carcinogenesis models of HCC such as, DEN-induced HCC are highly warranted in order to substantiate the therapeutic effects of Utt-B in preventing cancer initiation. Taken together, the findings of the current study highlight the anti-carcinogenic, anti-mutagenic and immunomodulatory effects of Utt-B, a US FDA-designated ‘Orphan drug’ against HCC.

## Author Contributions

Study concept and design: RJA. Acquisition, analysis, and interpretation of the data: MS, CKK, RPT, SCS, SS, KK, SUA, LRN, JS, SSV, AV, SJ, AJS. Drafting of the paper: SM, CKK, RJA Funding acquisition: RJA, KBH, KK; Review and editing of the manuscript: NPA, NI, RSL, KBH, RJA. All authors have read and agreed to the published version of the manuscript.

## Ethical Declaration

### Animal Ethical Clearance

Studies involving experiments with animals were conducted in accordance with ARRIVE guidelines and institutional guidelines under the approval from Institutional Animal Ethics Committee, Rajiv Gandhi Centre for Biotechnology. (CPCSEA Number: 326/GO/ReBiBt/S/2001/CPCSEA).

## Acknowledgements

We acknowledge the immense help provided Animal Research and Central Instrumentation facilities of Rajiv Gandhi Centre for Biotechnology, Thiruvananthapuram for the successful completion of the experiments. We also thank Mr. Dileep R K, Mr. Anwar, and Ms. Viji S for the technical help rendered during the animal experiments. We thank the service provider Redcliffe technologies, Bangalore for the successful completion of the transcriptomic profiling experiment.

## Standard Abbreviations

AFB1: Aflatoxin B1;
AFP: Alpha Fetoprotein;
Utt-B: Uttroside B;
HCC: Hepatocellular carcinoma;
H&E staining: Haematoxylin and Eosin staining;
H: hour;
LFT: Liver Function Test;
RFT: Renal Function Test;
PARP: Poly adenosine diphosphate-ribose polymerase;
ki67: Marker of proliferation;
PCNA: Proliferating cell nuclear antigen;
PBS: Phosphate buffered saline;
IP: Intra-peritoneal;
GAPDH: Glyceraldehyde 3-phosphate dehydrogenase;
FBS: Fetal bovine serum,
TNF-α: Tumor necrosis factor alpha;
IFN-ꝩ: Interferon gamma;
MCP-1: Monocyte Chemoattractant protein-1;
SCD-1: Stearoyl CoA desaturase;
SREBP 1-c: sterol regulatory element binding transcription factor;
TME: Tumor microenvironment,
DC: dendritic cells;
ICD: Immunogenic cell death.

---

**Supplementary Figure 1:**
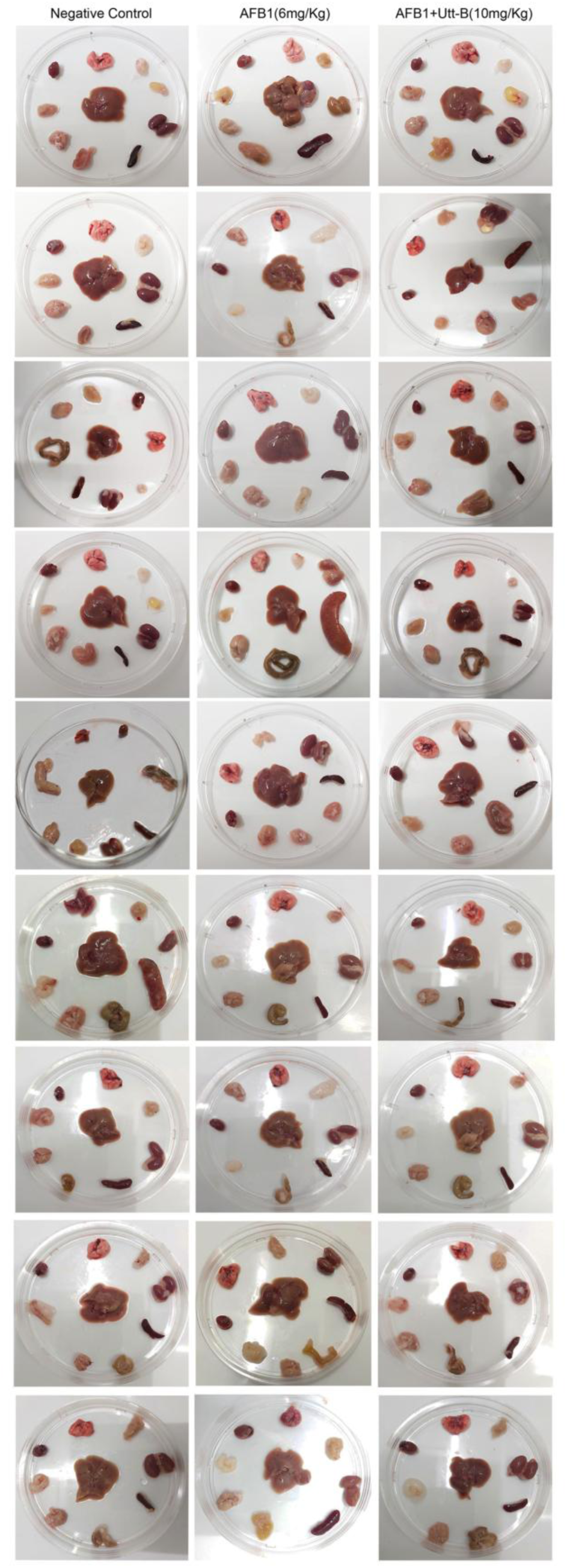
Representative photographs of organs of animals collected from different groups of AFB1-induced liver carcinogenesis model.

**Supplementary Figure 2:**
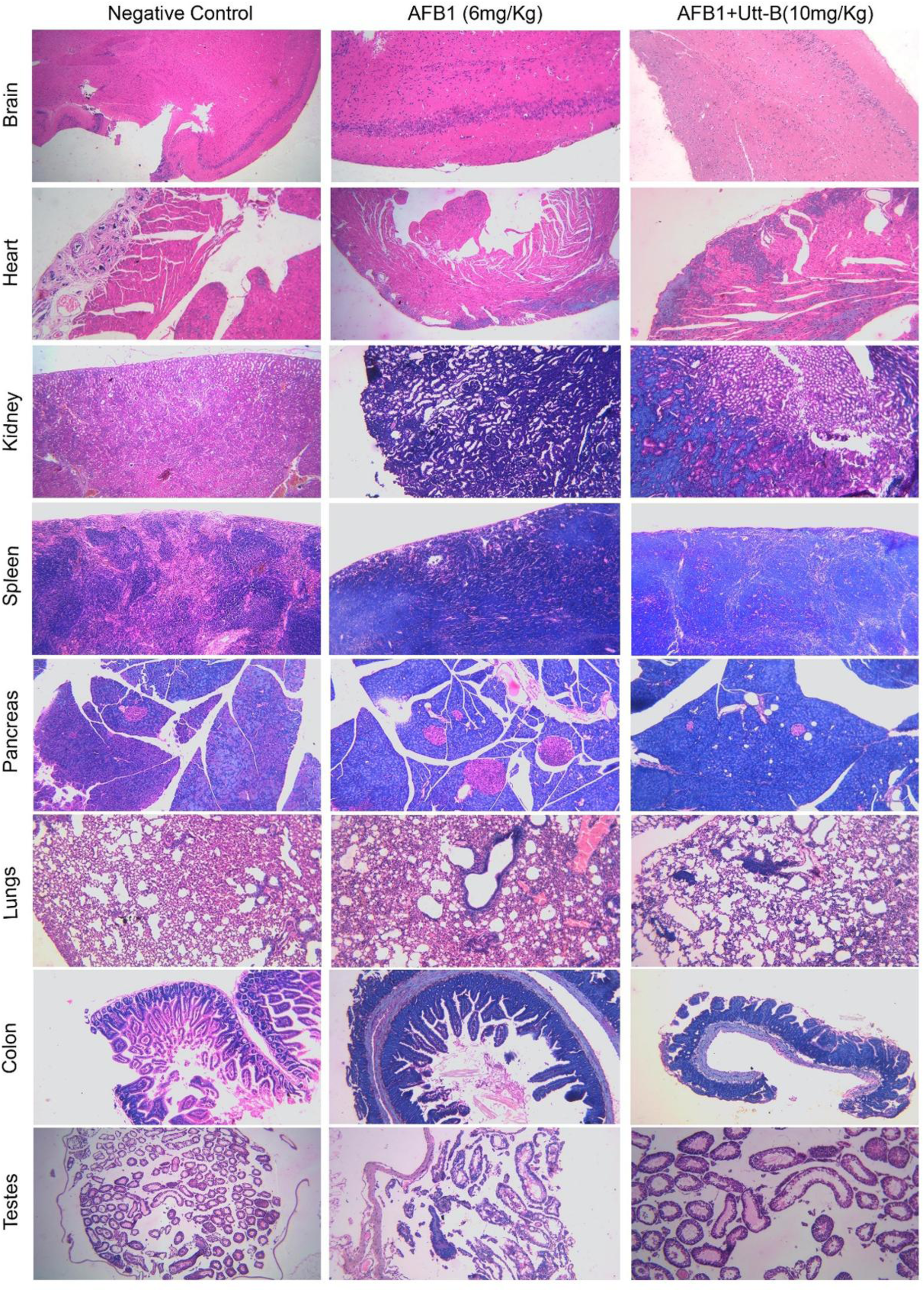
Histopathological evaluation of organs isolated from different treatment groups of AFB1-induced liver carcinogenesis model.

**Supplementary Table 1:**
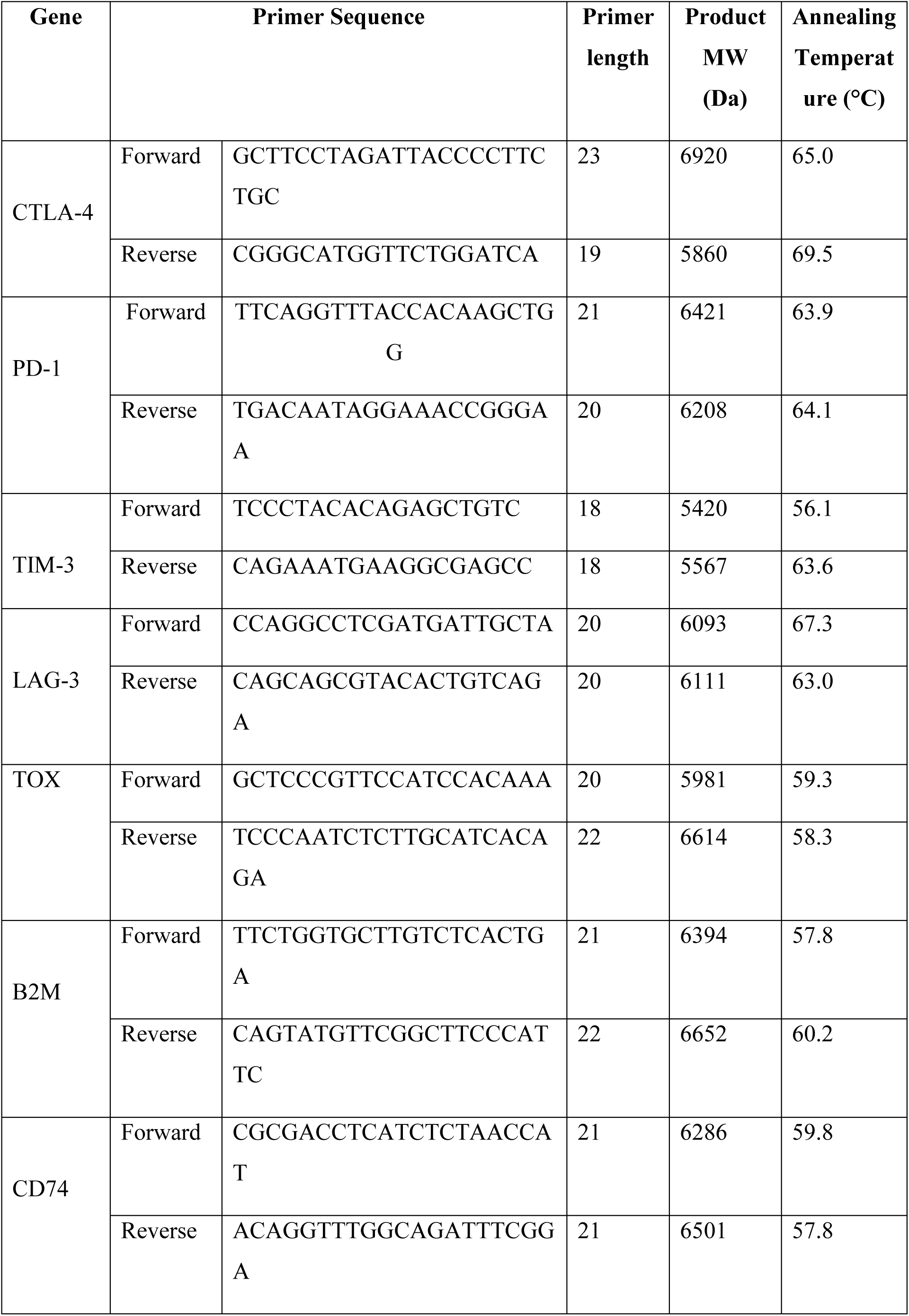
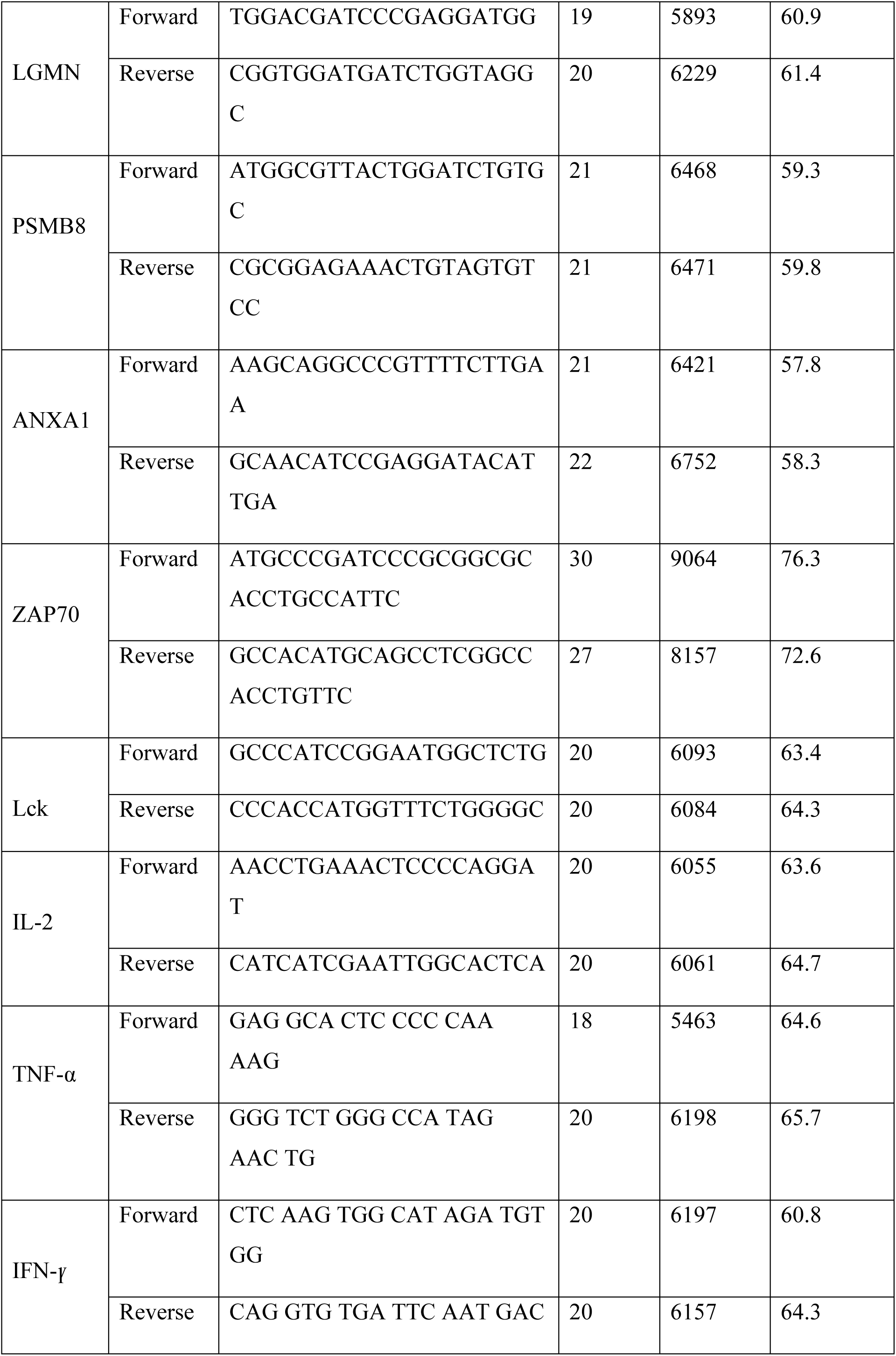
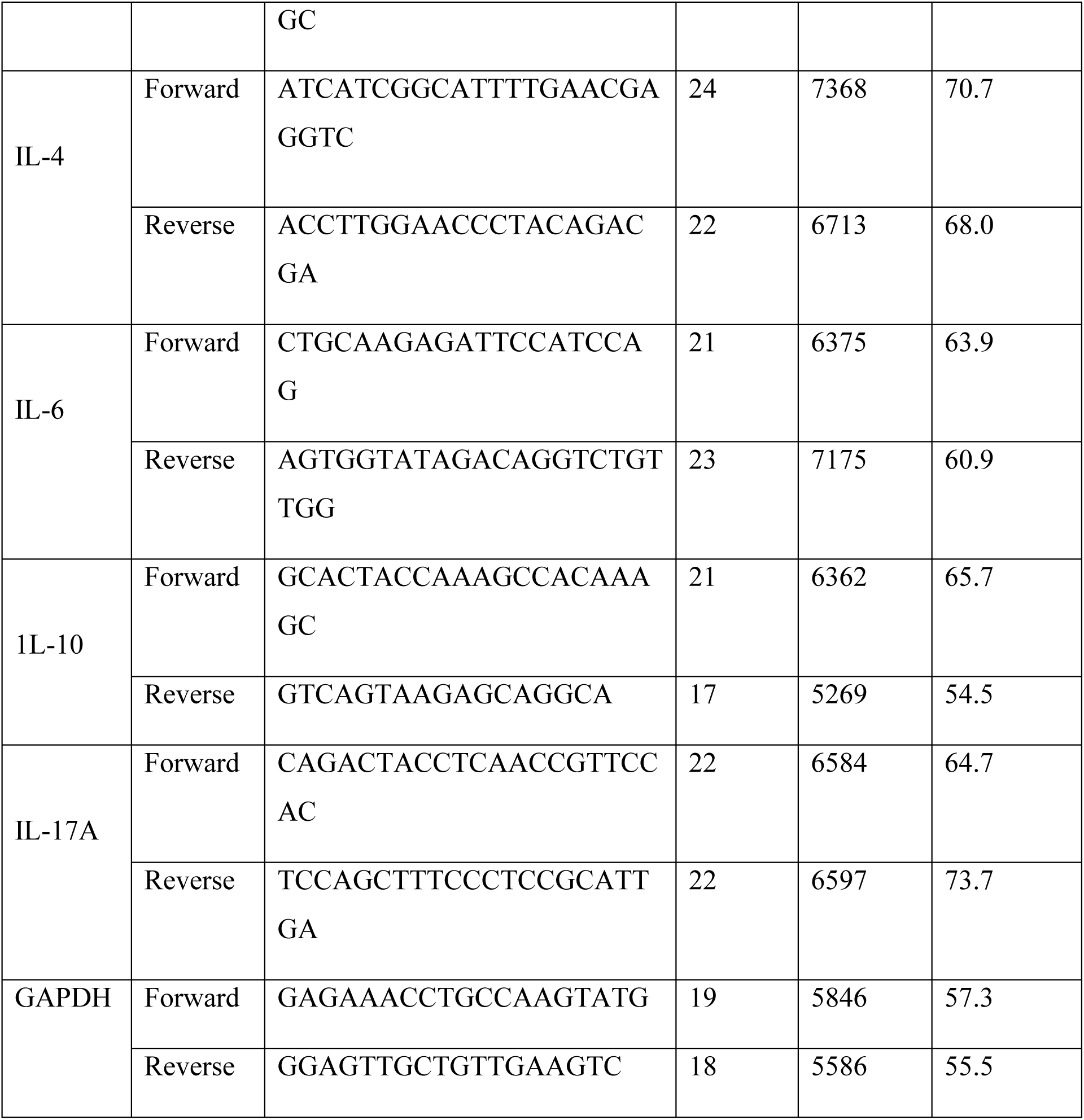
Details of Mouse Primers used in this study.

**Supplementary Table 2:**
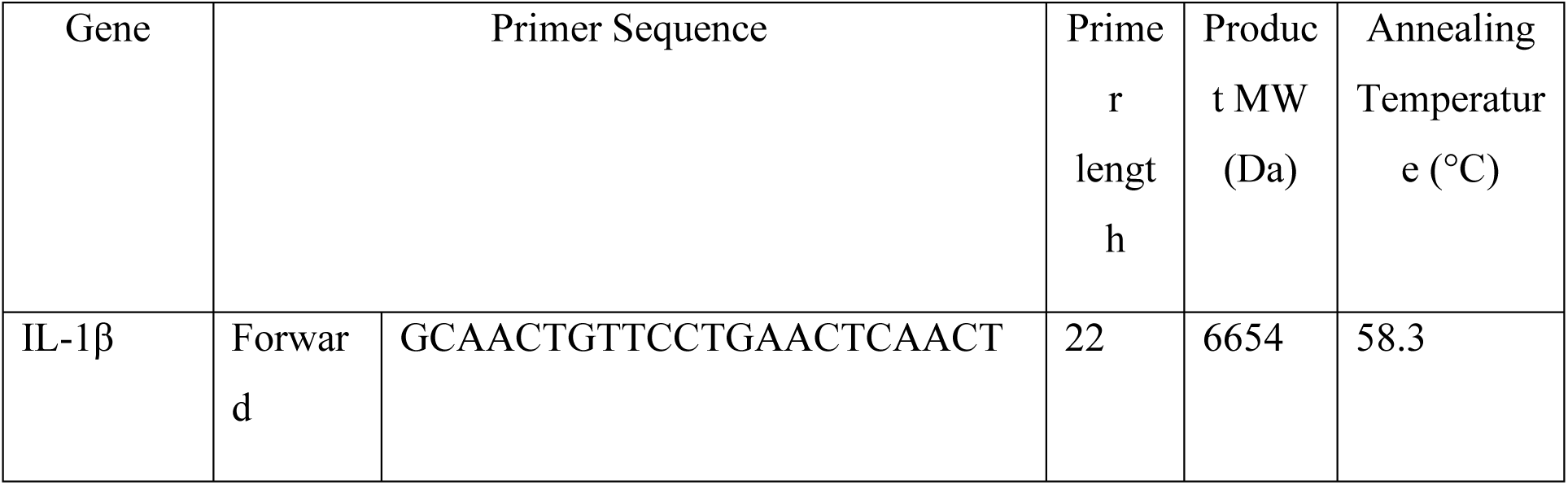
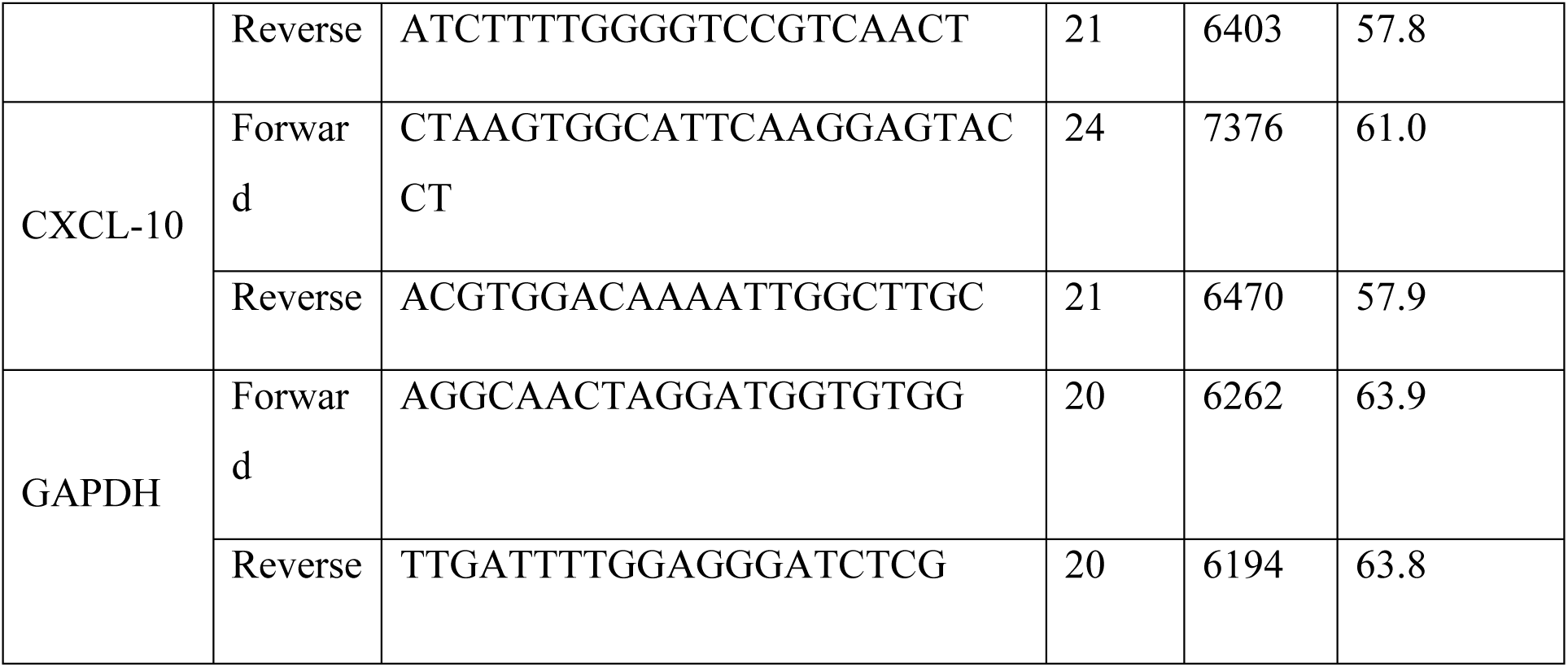
Details of Human Primers used in this study.

**Supplementary Table 3:**
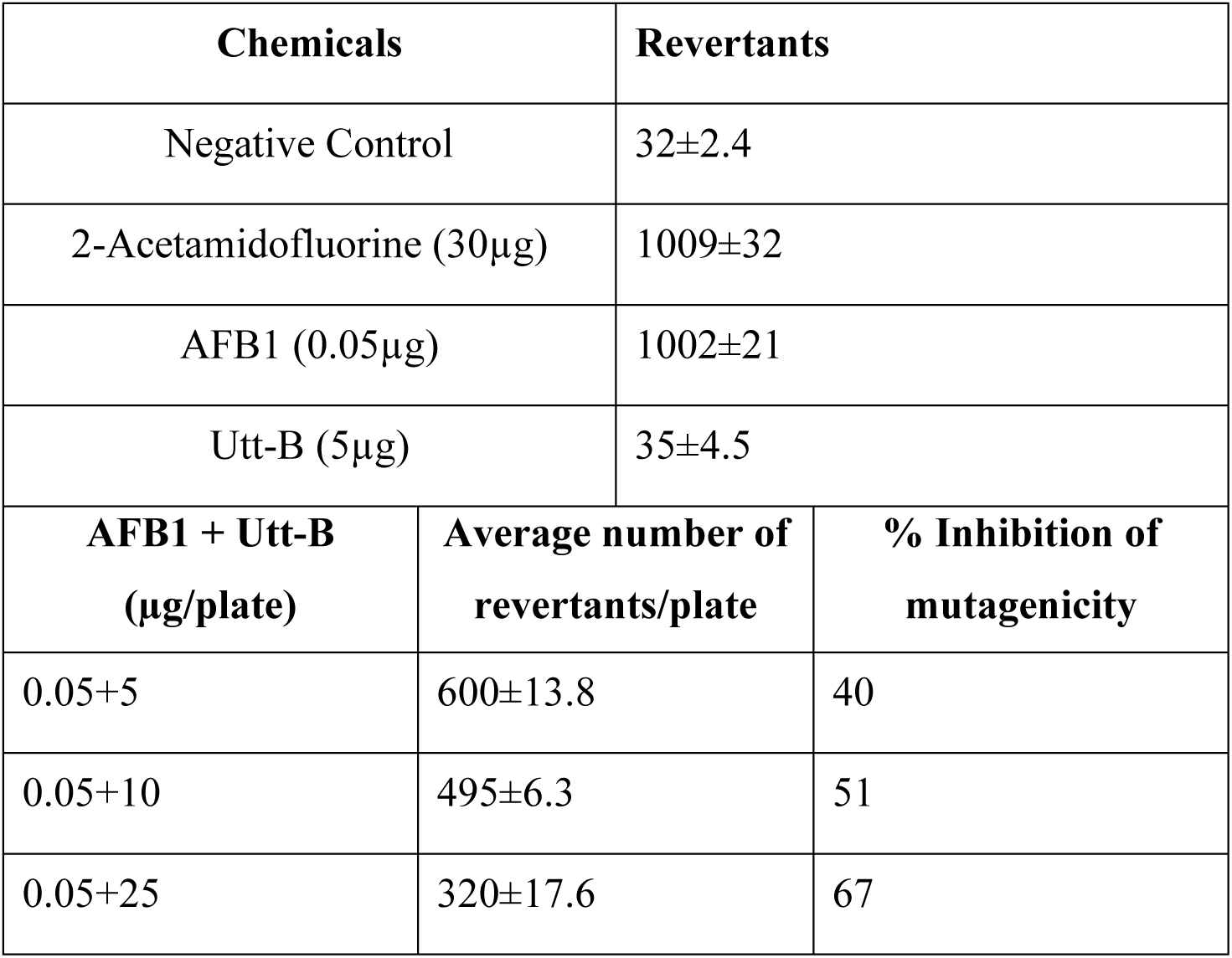
Utt-B inhibits AfB1-induced mutagenesis in TA98. The number of colonies in the control and experimental plates were counted and presented in the table (*P < 0.05)

| Chemicals |  | Revertants |
| --- | --- | --- |
| Negative Control |  | 32±2.4 |
| 2-Acetamidofluorine (30µg) |  | 1009±32 |
| AFB1 (0.05µg) |  | 1002±21 |
| Utt-B (5µg) |  | 35±4.5 |
| AFB1 + Utt-B<br>(µg/plate) | Average number of<br>revertants/plate | % Inhibition of<br>mutagenicity |
| 0.05+5 | 600±13.8 | 40 |
| 0.05+10 | 495±6.3 | 51 |
| 0.05+25 | 320±17.6 | 67 |

